# Pathway Association Studies Tool

**DOI:** 10.1101/691964

**Authors:** Adam Thrash, Juliet D. Tang, Mason DeOrnellis, Daniel G. Peterson, Marilyn L. Warburton

## Abstract

**Background:** In recent years, a bioinformatics method for interpreting GWAS data using metabolic pathway analysis has been developed and successfully used to find significant pathways and mechanisms explaining phenotypic traits of interest in plants. However, the many scripts implementing this method were not straightforward to use, had to be customized for each project, required user supervision, and took more than 24 hours to process data. PAST (Pathway Association Study Tool), a new implementation of this method, has been developed to address these concerns.

**Results:** PAST is implemented as a package for the R language. Two user-interfaces are provided; PAST can be run by loading the package in R and calling its methods, or by using an R Shiny guided user interface. In testing, PAST completed analyses in approximately one hour by processing data in parallel. PAST has many user-specified options for maximum customization. PAST produces the same results as the previously developed method.

**Conclusions:** In order to promote a powerful new method of pathway analysis that interprets GWAS data to find biological mechanisms associated with traits of interest, we developed a more accessible and user friendly tool. This tool is more efficient and requires less knowledge of programming languages to use than previous methods. Moreover, it produces similar results in significantly less time. These attributes make PAST accessible to researchers interested in associating metabolic pathways with GWAS datasets to better understand the genetic architecture and mechanisms affecting phenotype.

## Background

Genome-wide association study (GWAS) of complex traits in maize and other crops has become very popular to identify regions of the genome that influence these traits [1, 2, 3]. In general, hundreds of thousands of single nucleotide polymorphisms (SNPs) markers are each tested using F statistics for association with the trait, which assigns a *p*-value for the SNP-trait association. Individual marker-trait associations that meet the threshold set for the false discovery rate (FDR, the proportion of false positives among all significant results for some level α) are then studied in more detail to uncover hints as to the genetic architecture of the trait, and how best to improve it in the future. Many true associations may be missed in GWAS, however, because the threshold for FDR could be as low as α divided by the total number of SNPs being tested. In complex, polygenic traits, the effects of genes that exert only small effects on a trait may not meet the FDR threshold, especially if the effect value of the association is influenced by the environment. Additionally, alleles of many genes may be expressed only in specific genetic backgrounds and will only be useful when found in combination with the positive alleles of other genes in the same pathway [3]. These allelic combinations may not exist in the limited number of individuals in the GWAS panel. Thus, the statistical power of GWAS for detecting genes of small effect is limited by the strict levels set for FDR and by insufficient numbers of high-frequency polymorphisms found in most panels.

Metabolic pathway analysis focuses on the combined effects of many genes that are grouped according to their shared biological function [4, 5, 6]. This is a promising approach that can complement GWAS to give clues to the genetic basis of a trait. Originally developed to study differences in gene expression data in human disease studies [7], pathway analysis and association mapping have been used in medical research to find biological insights missed when focusing on only one or a few genes that have highly significant associations with a trait of interest [8, 9, 5, 10]. Pathway analysis has only just begun to be used as well in animal studies [11, 12]. In addition, biologically relevant pathways can be used to guide interpretation of large data sets produced by other high-throughput approaches like RNA sequencing, proteomics, and metabolomics.

More recently, GWAS-based metabolic pathway analysis has been used as a discovery tool to investigate the genetic basis of complex traits in plants. A pathway-based approach was used to study aflatoxin accumulation [13], corn ear worm resistance [14] and oil biosynthesis [15] in maize. Combining GWAS analysis with metabolic pathway analysis considers all genetic sequences positively associated with the trait of interest, regardless of magnitude, and jointly may highlight which sequences lead to mechanisms for crop improvement and which warrant further study and manipulation, for example, by gene editing. While combined GWAS and pathway analyses were highly successful in uncovering associated pathways, the analyses were slow and cumbersome, as the analysis tools were written in a combination of R, Perl, and Bash, and the output of each analysis was manually input into the next analysis. A single, unified and user friendly tool to accomplish this pathway analysis was lacking.

The Pathway Association Study Tool (PAST) was developed to facilitate easier and more efficient GWAS-based metabolic pathway analysis. PAST was designed for use with maize but is usable for other species as well. It tracks all SNP marker - trait associations, regardless of significance or magnitude. PAST groups SNPs into linkage blocks based on linkage disequilibrium (LD) data and identifies a tagSNP from each block. PAST then identifies genes within a user-defined distance of the tagSNPs, and transfers the attributes of the tagSNP to the gene(s), including the allele effect, R_2_ and *p*-value of the original SNP-trait association found from the GWAS analysis. Finally, PAST uses the gene effect values to calculate an enrichment score (ES) and *p*-value for each pathway. PAST is easy to use as an online tool, standalone R script, or as a downloadable R Shiny application. It uses as input TASSEL [16] files that are generated as output from the General Linear or Mixed Linear Models (GLM and MLM), or files from any association analysis that has been similarly formatted, as well as genome annotations in GFF format, and a metabolic pathways file.

## Implementation

PAST is implemented as an R package and is available through Bioconductor 3.9. PAST is based on a method developed by our research group [6]. The original method was subsequently used in two other maize studies [14, 15], but required users to customize Perl and R scripts and run BASH scripts. PAST’s implementation is completely in R and requires a user to install the package without needing to edit the source code. Two graphical user interfaces are available in the form of R Shiny applications. A generic version is available on CyVerse and Github, while a maize-specific version can be run on MaizeGDB [17] (explained below).

PAST processes data through four main steps. First, GWAS output data is loaded into PAST. This data comes in the form of statistics that reflect the effects of specific loci (e.g., SNPs) with a trait(s) of interest and LD data between loci. The SNPs are then associated with genes based on the allelic effects, *p*-values, and genomic distance between SNPs and genes. The genes and their effects are used to find significant pathways and calculate a running ES. Finally, the genes in these pathways are plotted to show a running ES for each pathway. A flowchart in Figure 1 shows the process.

**Figure 1:**
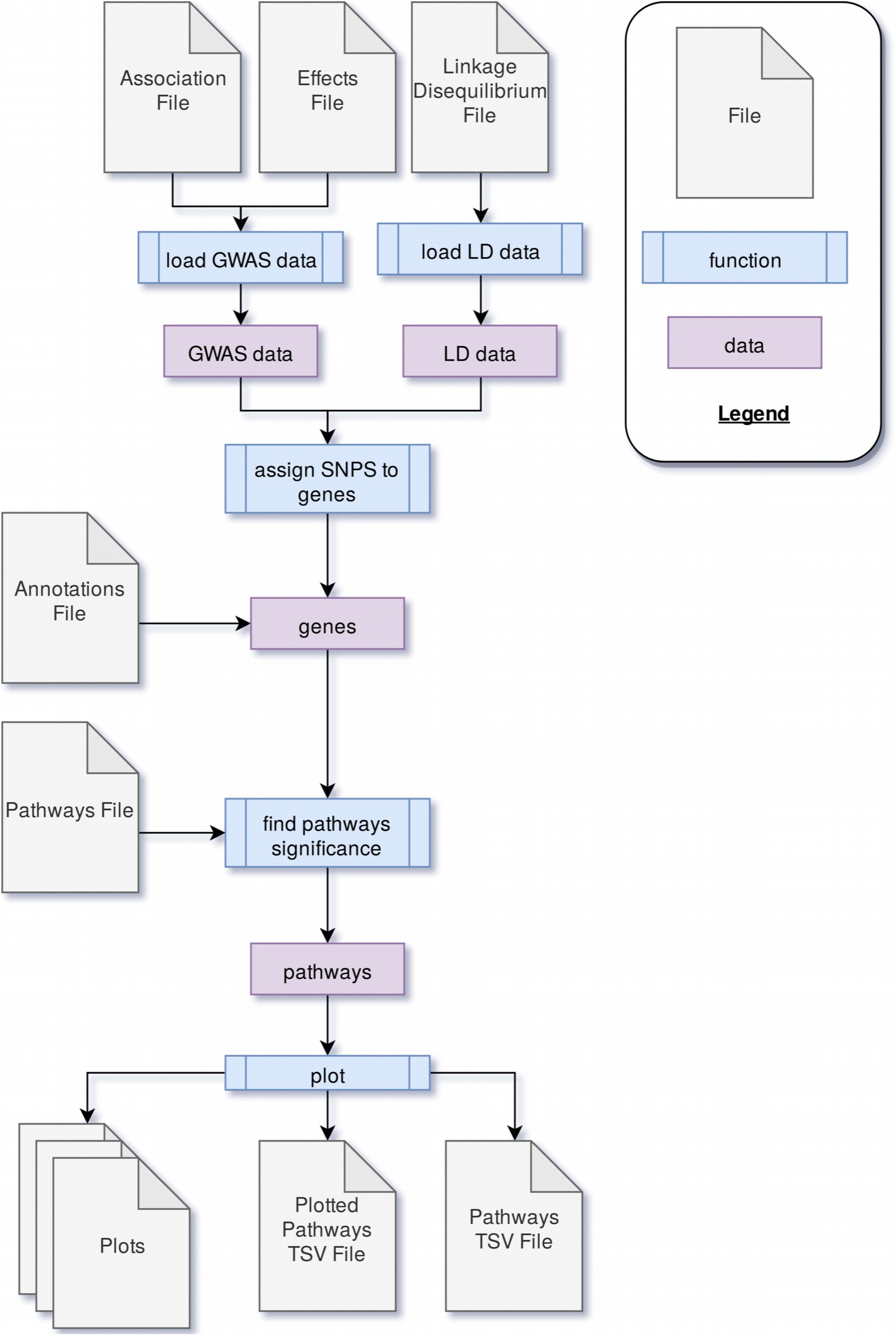
The process used in PAST.

### Loading Data

During the process of loading data, the GWAS data is filtered to account for any non-biallelic data. Any data with more or fewer than two alleles associated with that marker is discarded. Data without an R^2^ value (coefficient of determination the SNP/trait association) is removed as well, since later calculations rely on the R^2^ value. The effects data is associated with the statistics data in order to collect all data about a marker into a single dataframe.

The LD data is filtered to drop rows where the loci are not the same, and then extra columns are dropped. Only data about the locus, the positions, the sites, the distance between the sites, and the r^2^ value (coefficient of determination for LD) is retained. The remaining data is split into groups based on the locus.

### Assigning Genes

Genes are assigned the attributes of linked SNPs according to the method described in Tang *et al* [6]. SNPs are parsed into linked groups by identifying all pairs of SNPs with LD data that exceed a set cutoff r^2^ for linkage. SNP blocks occur when multiple SNPs are linked to one SNP in common. SNPs that are only linked to one other SNP are considered singly-linked SNPs, and SNPs not linked to any other SNPs are unlinked. In all cases, PAST follows an algorithm to identify one tagSNP to represent all linked SNPs in order to reduce the dimensionality of the dataset and identify which allele effect, *p* and R^2^ to transfer to the physically linked gene(s). Unlinked SNPs are by default identified as the tagSNP. For SNPs that are linked to a single other SNP, if both have the same effect sign (positive or negative), PAST identifies the one associated with the largest effect (absolute value) as the tagSNP. If the effects are equal, the second (more downstream) SNP is used. If, however, the effect signs are different, the SNP with the lowest *p*- value is used. If the *p*-values are the same and the signs are different, the SNP is labeled as problematic, since no assignment can be made, and no tagSNP is identified; these are dropped from the analysis.

The tagSNP within blocks of SNPs is identified by first counting the number of positive and negative effects in each linkage block. If the number of positive effects is greater, then the SNP with largest positive effect is chosen. If the number of negative effects is greater, then the SNP with the largest negative effect is noted. Ties between the number of negative and positive effects are broken by checking the sign of the SNP in common defining the block. The tagSNP is then the one with the largest effect and the same sign, and it is marked to indicate the number of SNPs in the block. Once all blocks have been reduced to a single tagSNP, the tagSNP is used to locate the nearby gene(s).

Once tagSNPs have been identified, the annotation files are checked to look for genes within a physical distance window provided by the user. The effect and the *p*-value of the tagSNP is transferred to the gene. The SNP-gene assignments are grouped by gene name, and if more than one SNP block or unlinked SNP was found to be linked to the same gene, each gene is tagged by counting the number of negative effect and the number of positive effect associations in the blocks linked to the same gene. If there are more negative effects, the most negative effect and *p*-value is assigned to the gene. If there are more positive effects, the positive effect and *p*-value is assigned to the gene. If there are more than one equally positive or equally negative effects, the effect with the lowest *p*-value is chosen and assigned to the gene. If there are an equal number of negative and positive effects, the effect with the greatest absolute value is selected. The number of linked SNPs is set to the total number of SNPs (SNPs within blocks plus blocks within genes) linked to that gene. Once all of the blocks of genes have been processed, the effects of each gene are used to find significant pathways.

### Finding Significant Metabolic Pathways

Significant pathways are found by using a previously described method [4, 6, 7]. User-input determines the minimum number of genes that a pathway must contain to be retained for processing (to avoid small sample size bias), the number of times the effects data are randomly sampled with replacement to generate a null distribution of ES, and the pathways database that is being used.

For each gene effects column (observed and randomly sampled), the effects are sorted and ranked from best to worst (and whether this is in increasing or decreasing order depends on the trait under study). The ES running sum statistic increases for genes in the pathway and decreases for genes not in the pathway. The amount of increase for genes in the pathway is weighted by the absolute value of the effect. The pathway ES is the largest positive value calculated for the running sum statistic.

Pathway significance is determined by comparing the observed ES with the ES for the null distribution. The mean and standard deviation for the null distribution are used to normalize the observed ES so that *z* scores can be obtained. *P*-values are computed from the *z* scores using the (1-pnorm) function. Since multiple hypothesis testing is still a concern, an FDR-adjusted *p*-value (known as q-value) is calculated using the qvalue package in R [18].

### Plotting

Based on user input, the pathways can be filtered for significance (either *p*-value or q-value), or the top *n* pathways can be selected. Rugplots for each pathway in the set of significant pathways are plotted as the last step. The x-axis shows the rank of each gene effect value; the y-axis shows the value of the ES running sum statistic as each consecutive gene effect value is processed. An x-intercept line indicates the highest point of the ES. Small hatch marks at the top of the image indicate the position of the effect of all genes in the pathway. An example rugplot is provided in Figure 2.

**Figure 2:**
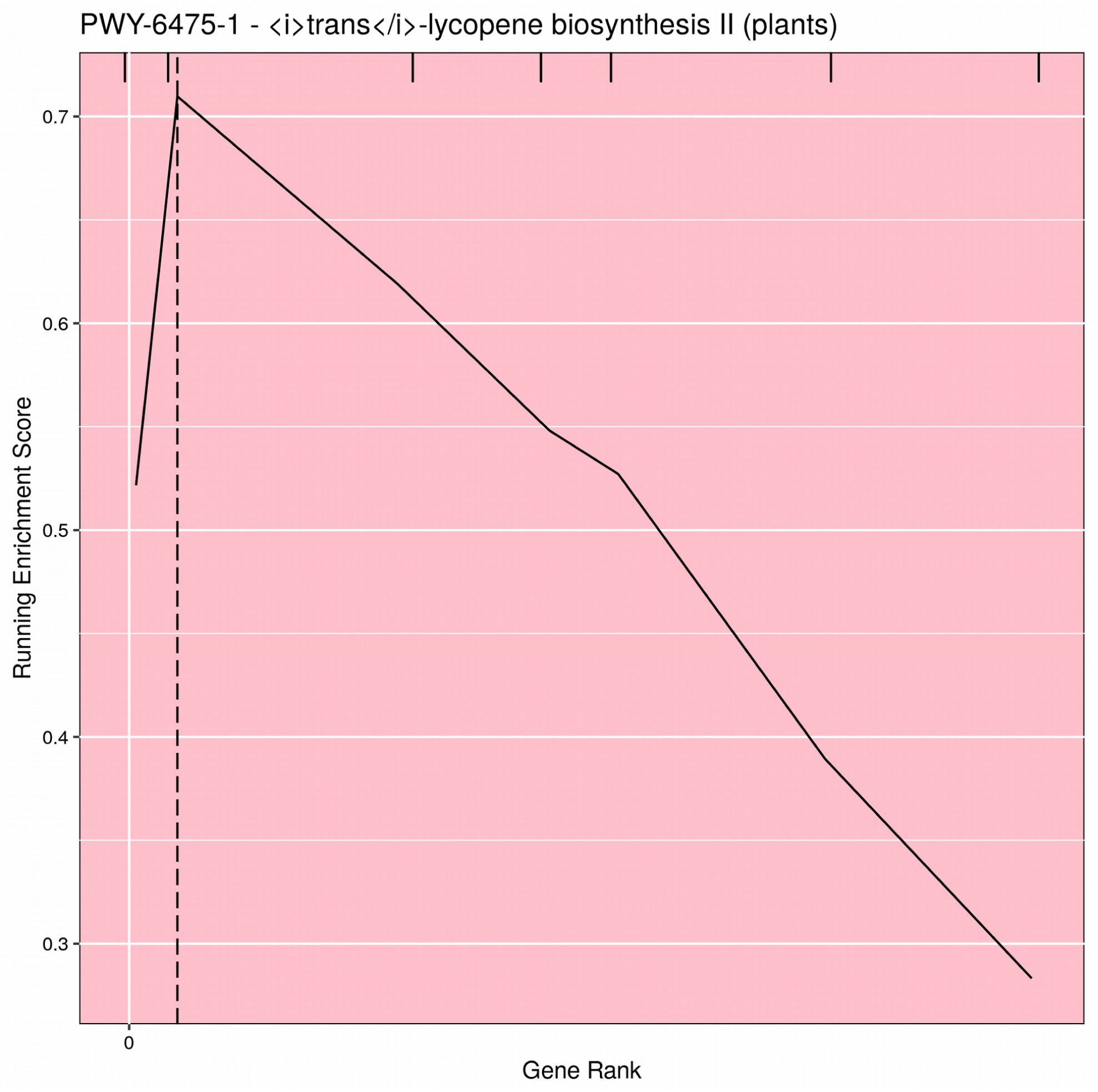
Example PAST Output.

### R Shiny Applications

Two versions of an R Shiny application that use PAST have been developed. These R Shiny applications provides a guided user interface that sets analysis parameters in PAST; they can also upload a saved set of results to explore again. The version available on Github and CyVerse allows a user to run a new analysis by selecting their data, annotations, and pathways depending on the species being studied. The version available on MaizeGDB [17] allows a user to upload their data and select specific versions of the maize annotation and pathways databases available on MaizeGDB. A screenshot of the generic R Shiny application is provided in Figure 3.

**Figure 3:**
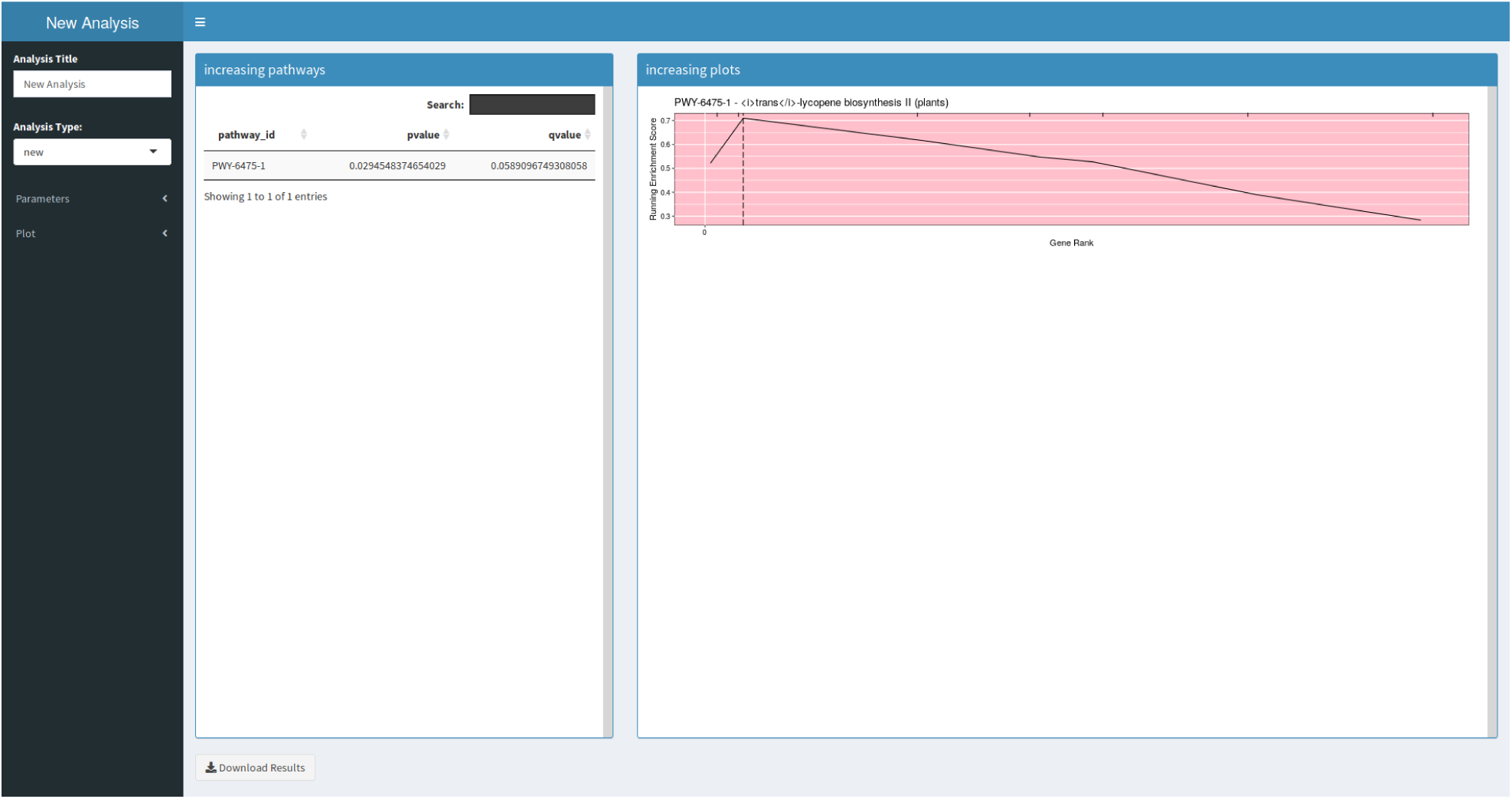
A screenshot of the R Shiny application.

## Results and Discussion

PAST is run by calling its functions with GWAS data from within an R script or by using an included R Shiny interface. PAST will allow a new interpretation of GWAS results, which should identify associated pathways either when one or a few genes are highly associated with the trait (these would have been identified by the GWAS analysis directly); or when many genes in the pathway are moderately associated with the trait (these would not necessarily have been identified by the GWAS analysis). Such an interpretation will add both additional results, and biological meaning to the association data, as was seen with oil biosynthesis in studies by Li et al. [19, 15]. While PAST may be useful in bringing biologically useful insights to a GWAS analysis, it will not be able to find order from a chaotic dataset if environmental variation, experimental error, or improper analysis models were used in the association analysis. For strong data sets, however, it may find pathways where GWAS found few or no significant associations which, taken in isolation, shed no real light on the genetic mechanisms underlying the traits under study. PAST may be able to overcome this limitation, and may in addition be able to identify epistatic interactions between genes in the same pathway [20].

PAST was tested using data from three previous corn GWAS on kernel color (261147 SNPs), aflatoxin resistance (261184 SNPs), and total oil production (525107 SNPs). Kernel color and aflatoxin analyses were run on a desktop computer with 32GB of memory, a 4GHz Intel Core i7 with four processors, and solid-state storage. All four processors were used when testing PAST. The kernel color test completed in 63 minutes; the aflatoxin test completed in 62 minutes. The total oil test was run on a system with the following specifications: Intel(R) Xeon(R) CPU E5-2680 v2 at 2.80 GHz with 64 GB of RAM connected to a lustre v.2.5.3 file system via an FDR InfiniBand interconnect. The analysis of the total oil trait completed in 71 minutes. Using the previous method, these analyses took 24 hours or more, depending on how attentive the user was to starting the next step in the process. The results of the analyses of all three traits were comparable when generated with PAST or with the previous method.

The use of metabolic pathway analysis to derive functional meaning from GWAS results has been used extensively in human disease studies, and methodologies and tools similar to PAST have been published for use with annotated human pathways. Some methodologies reviewed by Kwak and Pan [21] include GATES-Simes, HYST, and MAGMA. Two tools for human GWAS pathway studies have been published, including GSA-SNP2 [10] and *Pascal* (a Pathway scoring algorithm) [22]. However, these tools would need to be extensively modified to work with any set of user-supplied pathways. Some of the analysis procedures, however, should be considered in future versions of PAST. An analysis with PAST should be illuminating for any outcrossing plant species and any animal and human datasets for which annotated pathway/genome databases (or related model organism databases) and GFF annotations are available. However, for inbreeding and polyploid species, the assignment of SNPs to genes may be complicated by very long LD blocks which may contain multiple, equally linked genes, or homology to more than one genome. Tests will be run to see if these problems can be overcome.

## Conclusions

In conclusion, we present PAST, a tool designed to use GWAS data to perform pathway analysis. PAST is faster and more user-friendly than previous methods, requires minimal knowledge of programming languages, and is publicly available at Github, Bioconductor, CyVerse and MaizeGDB.

## Availability and requirements

Project name: PAST

Project home page: https://github.com/IGBB/PAST/

Shiny apps: https://apps.maizegdb.org/PAST/, https://apps.maizegdb.org/maizeGDB_PAST/, <CyVerse link once we have it>

Bioconductor: https://doi.org/doi:10.18129/B9.bioc.PAST

Operating system(s): Platform independent

Programming language: R 3.6

Other requirements: R packages dplyr, rlang, iterators, parallel, foreach, doParallel, qvalue, rtracklayer, ggplot2, shiny, shinydashboard, gridExtra

License: GNU GPLv3

Any restrictions to use by non-academics: None

## List of abbreviations

(PAST): Pathway analysis study tool
(GWAS): genome-wide association study
(LD): linkage disequilibrium data
(SNP): single nucleotide polymorphism
(ES): enrichment score

## Declarations

All manuscripts must contain the following sections under the heading ‘Declarations’:

### Ethics approval and consent to participate

NA

### Consent for publication

NA

### Availability of data and materials

The example data used is available on the project’s GitHub, in the example folder.

## Authors’ contributions

AT contributed to the development of the project, directed development of the software, and contributed to the manuscript. JT developed the method in its original implementation, advised development of PAST, and contributed to the manuscript. MD contributed to the development of PAST, and contributed to the manuscript. DGP advised development of the project. MW provided example data, advised development, contributed to the manuscript, and directed the entire project. All authors read and edited the manuscript.

## Acknowledgements

The authors gratefully acknowledge helpful suggestions from Dr. Andy Perkins and Mr. Mark Arick II; and Drs. Peter Bradbury, Brandon Monier, and Mr. Terry Casstevens for their thoughtful manuscript review and suggestions.

